# Developmentally programmed epigenome regulates cellular plasticity at the parental-to-zygote transition

**DOI:** 10.1101/2022.03.01.482564

**Authors:** Ryan J. Gleason, Christopher S. Semancik, Gitanjali Lakshminarayanan, Xin Chen

## Abstract

During metazoan development, the dramatic potency change from germline to embryos raises an important question regarding how the new life cycle is reset. Here, we report a tightly regulated epigenome landscape change from the parental germline to embryos in *C. elegans*. The epigenome is enriched with histone H3 in early-stage germ cells but switches to a histone variant H3.3-enriched epigenome in the mature egg. This H3.3-dominant epigenome persists in early-stage embryos until gastrulation, when the epigenome becomes H3 abundant again. We further demonstrate that this developmentally programmed H3 → H3.3 → H3 epigenome landscape change is regulated through differential expression of distinct histone gene clusters and is required for both germline integrity and early embryonic cellular plasticity. Together, this study reveals that a bimodal expression of H3 *versus* H3.3 is important for epigenetic reprogramming during gametogenesis and embryonic plasticity.

**One Sentence Summary:** Developmentally programmed epigenome resets cellular plasticity at the parental-to-zygote transition in *C. elegans*.

## Main text

A fundamental and important developmental biology question is how a fertilized egg, upon division and differentiation, becomes a highly organized embryo. On the one hand, gametogenesis represents one of the most dramatic terminal cellular differentiation pathways. On the other hand, upon fertilization, the totipotency is regained in the zygote (*1*). During these processes, epigenetic information is robustly and extensively established, erased, and reset (*2-4*). Both canonical histones and histone variants play important roles in epigenetic regulation. The canonical, or replication-coupled (RC) histones (i.e., H3, H4, H2A, H2B) are expressed during S-phase and mainly incorporated into the genome during DNA replication. Histone variants are typically replication-independent (RI), expressed throughout the cell cycle, and regulate a variety of biological processes (*5-7*). Intriguingly, unicellular organisms such as yeast only have H3.3-like histones, whereas metazoans have both H3.3-like and H3-like histones, suggesting that H3 could be responsible for more specialized roles in metazoan development (*8, 9*), such as regulating distinct cell fates.

Metazoan RC histones are typically encoded by gene clusters found at multiple chromosomal locations (*10*). For example, the human genome has 14 histone *H3* genes on two chromosomes, while the *C. elegans* genome has 15 histone *H3* genes on four chromosomes (*11*). Due to the high sequence similarity among the multiple *H3* genomic loci, defining contributions of individual histone *H3* genes has remained a challenge due to the lack of precise genetic tools. Furthermore, H3 and the histone variant H3.3 share a 97% similarity among their amino acid sequences, and are two of the most conserved proteins among all eukaryotic organisms (*8, 12*). Despite the sequence and structural similarity between H3 and H3.3 (*13*), they exhibit distinct interactions with histone chaperones and have different genomic distributions (*6, 14, 15*). Functionally, H3.3 is often associated with active transcription and enriched with post-translational modifications (PTMs) such as H3K36me2 and H3K4me3 (*16-20*). In contrast, PTMs associated with more repressive chromatin, such as H3K27me2/3 and H3K9me2/3, occur preferentially on H3 (*17-19*). Furthermore, recent studies in several organisms have suggested a conserved role for H3.3 during gametogenesis and early embryonic development in mice (*21-24*), *Drosophila* (*25-27*), and *X. Laevis* (*28*). In *C. elegans*, removal of H3.3 is not lethal, but reduces fertility and viability in response to stress (*29*). However, it is unknown whether and how individual *H3* genes act in *C. elegans* and how they interact with H3.3. Here, we characterized expression patterns and roles of individual histone *H3* genes, and compared them to the histone variant *H3.3*, during *C. elegans* gametogenesis and embryogenesis. We identified unique developmentally programmed expression and function of the 15 histone *H3* genes, which uniquely influence the epigenome in the germline and restrict embryonic plasticity by providing a barrier against cellular reprogramming.

To investigate the spatiotemporal dynamics of each *H3* gene cluster in the *C. elegans* genome, we used CRISPR/Cas9 genome engineering to systematically tag endogenous histone genes (*30*), leading to a series of histone *H3* knock-in lines that produce fusion proteins with different fluorescent tags such as Dendra2, mCherry, and eGFP, at their C-termini (*31*). Using quantitative, live cell imaging to characterize histone H3 incorporation patterns, we uncovered two major classes of *H3* genes based on their distinct expression patterns during gametogenesis and early embryogenesis. Surprisingly, only a small subset (4 of 15 *H3* genes, Class I) are expressed in both the germline and somatic lineages, which include *his-45, his-55, his-59*, and *his-63* (Fig. 1A and B, fig. S1A and B). The majority of *H3* genes (10 of 15, Class II) are not detectable in the germline but only in the somatic cell lineages, including members of the HIS1 (*his-2*), HIS2 (*his-6*), HIS3 (*his-9, his-13, his-25*, and *his-42*), HIS4 (*his-17, his-27* and *his-49*), and HIS5 (*his-32*) histone gene clusters (Fig. 1B and fig. S1C). Notably, expression patterns among different *H3* genes within the same gene cluster are consistent with each other. Of the 15 histone *H3* genes, *his-40* is unique considering it can only be detected in a subset of somatic lineages (Fig. S1D and table S1). Consistent with our results, recent RNA profiling assays have revealed two major RC histone gene clusters based on their distinct expression patterns. The Class I histone *H3* genes (*his-45, his-55, his-59*, and *his-63*) show significant expression at the earliest embryonic timepoints as well as from the gonad, suggesting their potential roles in intergenerational epigenetic inheritance (*32*). In contrast, the Class II histone *H3* genes show a largely zygotic expression pattern. The consistent patterns between histone H3 proteins and transcripts indicate that their expression is not subject to significant post-transcriptional regulation.

**Fig. 1.**
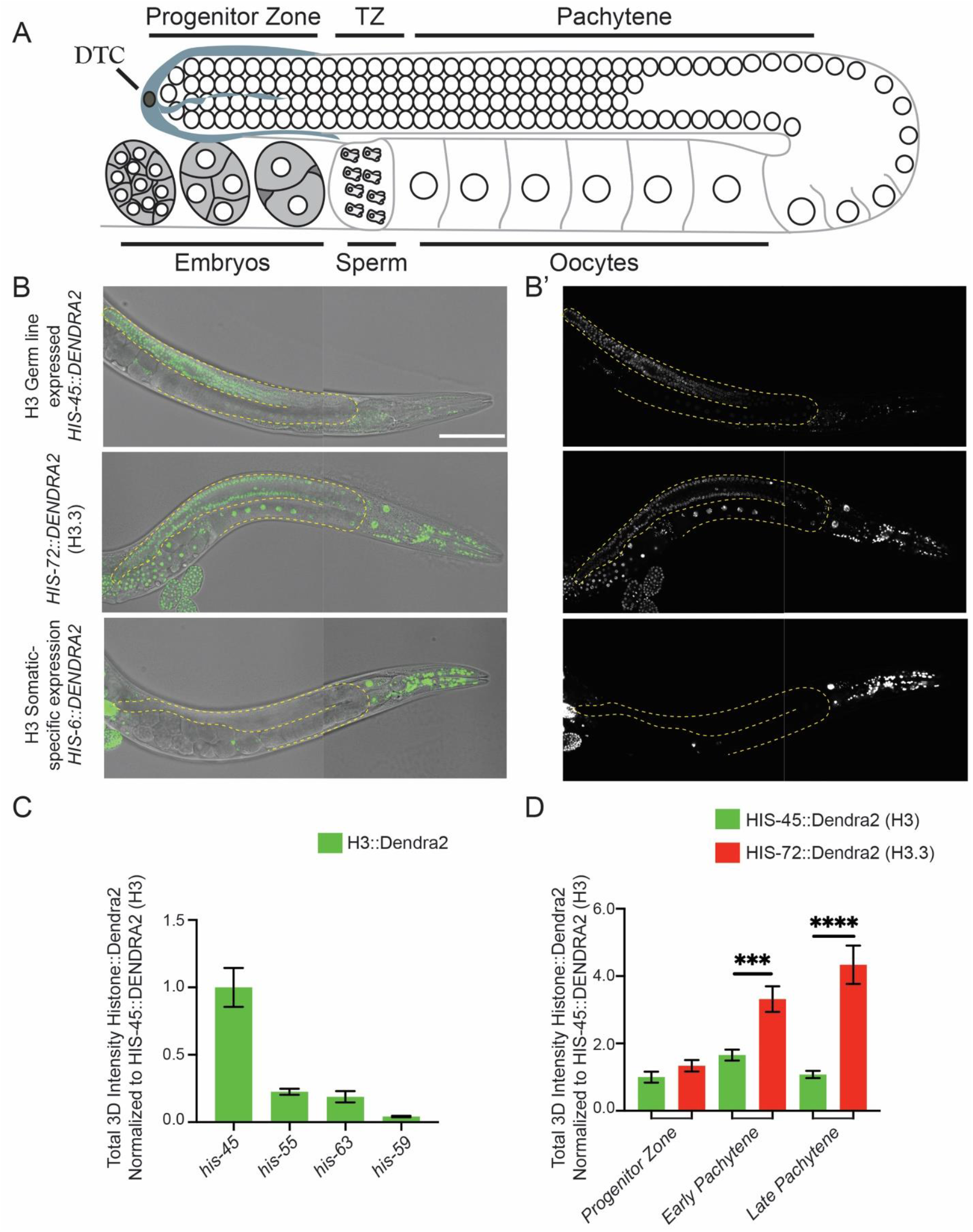
Differential incorporation of the replication-coupled histone *H3* gene family. (**A**) Illustration of the *C. elegans* hermaphrodite gonad. Germline nuclei are arranged in a spatiotemporal pattern progressing from the <u>D</u>istal <u>T</u>ip <u>C</u>ell (DTC) through the mitotic zone (Progenitor Zone), the <u>T</u>ransition <u>Z</u>one (TZ), to the Pachytene stage until the mature oocyte. Oocytes become fully cellularized by late diakinesis, pass through the spermatheca filled with sperm, and undergo early embryonic development *in utero*. (**B – B’**) Representative differential interference contrast (DIC) image and fluorescence micrographs of CRISPR-tagged HIS-45::DENDRA2 (H3) (Green), the dashed lines outline the gonads. HIS-45::DENDRA2 is an example of one of four Class I *H3* genes whose expression are detectable in both germline and somatic tissues. CRISPR-tagged HIS-72::DENDRA2 (H3.3), shows ubiquitous expression including all somatic tissues and throughout the germline. CRISPR-tagged *H3* gene HIS-6::DENDRA2 expression is not detectable in the germline, but is detectable throughout the somatic cell lineage, representing the expression pattern of 10 of 15 Class II histone *H3* genes. All images from (**B – B’**) are taken using the same imaging settings. (**C**) Quantification of all four *H3* genes expressed in the germline. For each condition, the total amount of H3::DENDRA2 present per mitotic/progenitor germline nuclei was quantified including *his-45* (n=9), *his-55* (n=15), *his-63* (n=15), and *his-59* (n=18). (**D**) Quantification of the most robust germline-expressed *his-45 H3* gene along the distal-proximal gonadal regions: <u>G</u>ermline <u>S</u>tem <u>C</u>ell (GSC) region (n=18), early pachytene (n=18), late pachytene (n=18) and *his-72 H3.3* gene: GSC region (n=21), early pachytene (n=21), late pachytene (n=21). All quantifications = average ± SE. *P*-value: unpaired t test **** *P*≤*0.0001*, *** *P*≤*0.001*. Scale bar: 100 μm.

In the *C. elegans* gonad, germ cells are organized in a temporally and spatially organized manner along the longitude axis (Fig. 1A). Examination of the four Class I *H3* genes revealed that *his-45* is the highest expressed in the germline (Fig. 1C and fig. S1B). In order to directly compare H3 *versus* H3.3 quantitatively in a spatiotemporally specific manner, we generated two CRISPR-tagged *H3.3* strains, *H3.3::Dendra2* and *H3.3::mCherry*, respectively (Fig. 1B and fig. S2A). We then used the *H3.3::Dendra2* for quantitative temporal imaging analyses to compare with *H3::Dendra2* strains. On the other hand, the *H3.3::mCherry* was used with different *H3::eGFP* strains for their potential distinct localization and chromosomal association. Based on these studies, we found that H3 and H3.3 display distinctly dynamic patterns: As germline nuclei move proximally, an approximate 4-fold increase of H3.3 was detected in late pachytene germ cells (Fig. 1D). In contrast, H3 is enriched in the distal mitotic region. In addition, the condensed state of chromosomes during meiotic prophase allows identification of individual chromosomes (Fig. S1A). Using antibodies against specific histone PTMs, H3K36me2 is enriched on autosomes while the X chromosome is deficient of H3K36me2 but is primarily associated with H3K27me2/3, consistent with the repressive chromatin status for the X chromosome (*33, 34*). In accordance with previous reports, H3.3 was found to accumulate on the autosomes but is depleted from the X chromosome, based on its anticorrelation with H3K27me2/3 and co-localization with H3K36me2 (*35-37*) (Fig. S2B). Contrastingly, germline-expressed H3 was enriched in H3.3-depleted regions, including the X chromosome, marked by its co-localization with H3K27me2/3 but anticorrelation with H3K36me2 (Fig. S2B). These results suggest that H3 carries unique modifications that establish and/or maintain repressed regions of the epigenome. Furthermore, we found both H3.3 and H3 are retained in mature sperm and oocyte chromatin (Fig. S2A), indicating that they may regulate the transition from germline to zygote.

In *C. elegans*, embryos initiate zygotic transcription in the blastomeres beginning at the four-cell stage, while the onset of gastrulation begins at the 26-cell stage (*38-40*). Interestingly, we observed a low level of H3 but a high level of H3.3 throughout the rapid cell cycles of early embryogenesis (Fig. 2A). However, when gastrulation initiates, a drastic increase of all 15 histone *H3* genes was detected (Fig. 2A-B). Consistent with the above results (Figure 1), the expression timing of the *H3* genes coincide with each other within the same class (i.e., Class I *versus* Class II), suggesting they are developmentally coordinated for co-expression. Using *his-45* (Class I) and *his-6* (Class II) to represent the two classes of *H3* genes, the *his-45* was detectable throughout embryogenesis, including the earliest cell divisions, and increased gradually upon gastrulation (Fig. 2A-B). Remarkably, expression of the soma-specific *his-6* gene was not detectable until the onset of gastrulation, and increased rapidly over the subsequent cell divisions (Fig. 2A-B). Quantifying expression of these two H3 classes highlights the global change in the epigenome, from an H3.3-enriched epigenetic landscape in late germline and early embryos to an H3-enriched epigenome in late embryos (Fig. 2A-C). Furthermore, this change is specific to the somatic lineage cells, as live cell imaging revealed that the P lineage for primordial germ cells does not display such an increased H3 incorporation throughout embryogenesis but remained enriched with H3.3 (Fig. 2C and movie S1).

**Fig. 2.**
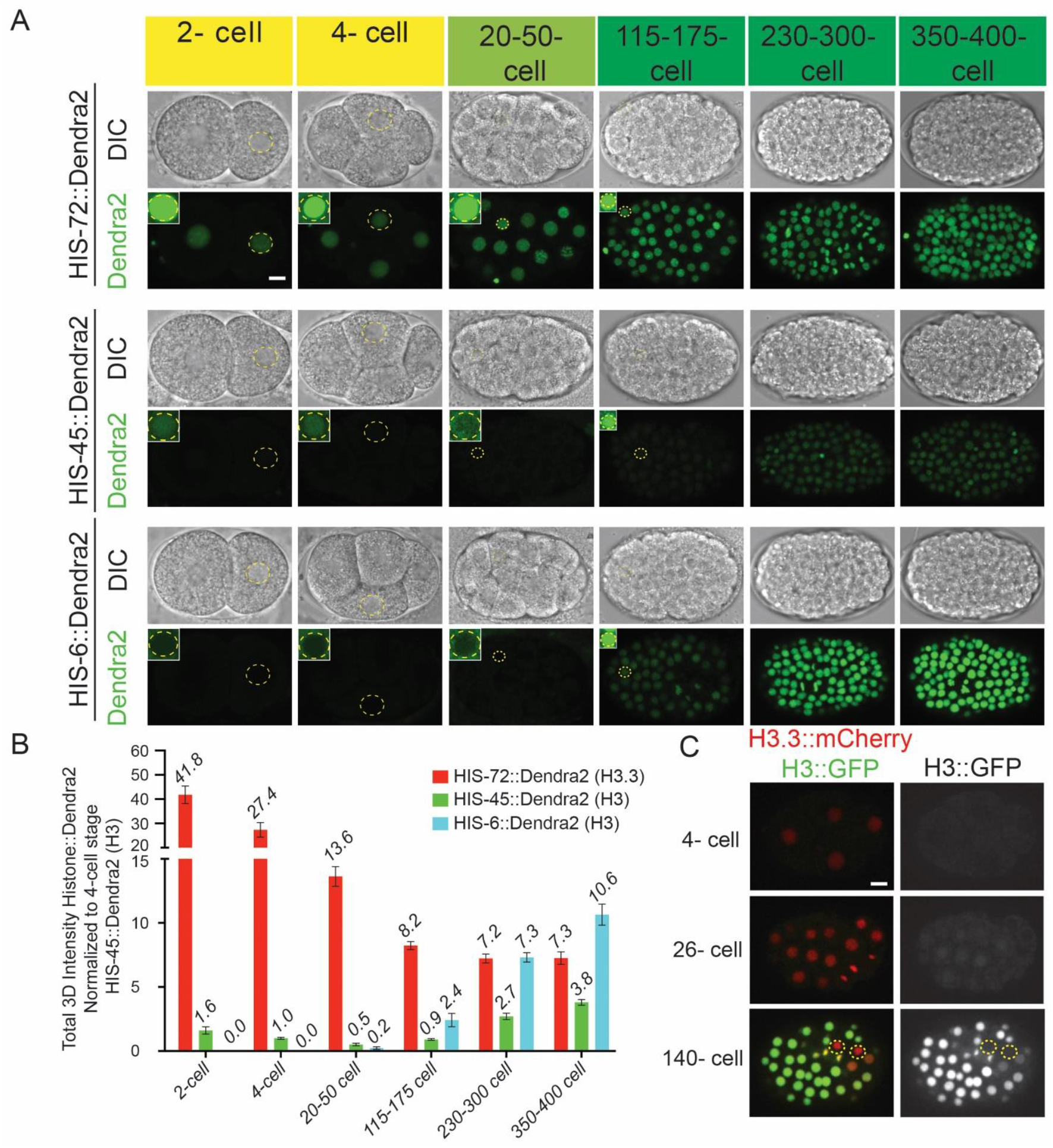
The early embryonic epigenomic landscape is developmentally programmed to switch from histone variant H3.3-enriched to histone H3-enriched upon gastrulation. (**A**) Representative images of embryos expressing H3.3-encoding *his-72::dendra2*, germline-expressed, H3-encoding *his-45::dendra2*, and somatic H3-encoding *his-6::dendra2* throughout the designated stages of embryogenesis including pre-gastrula (2- and 4-cell, yellow), gastrula (21-50 cell, light green), and after the onset of gastrulation (115 – 400 cell, green), DIC (top) and Dendra2 (bottom). All images from (**A**) are taken using the same imaging settings, insets demonstrate that *his-72* (H3.3) is readily detectable, whereas *his-45* (H3) is barely detectable by increasing the brightness; however, *his-6* (H3) is undetectable even with enhanced brightness but only becomes detectable at the onset of gastrulation (at the 26-cell stage). (**B**) Quantification of data sets from (**A**). For each embryonic stage, the total amount of DENDRA2 representing either H3 or H3.3 per nuclei were quantified. All quantifications = average ± SE. (**C**) Snapshots from time-lapse movies of embryos expressing HIS-72::mCherry (H3.3) and HIS-55::GFP (H3) (see movie S1). The dashed circles outline the primordial germ cells (PGCs). Scale bar: 5 μm.

We next asked whether the germline expressed Class I *H3* genes are required for normal gametogenesis and fertility. To address this question, we generated deletion alleles in two of the lowly expressed Class I *H3* genes, *his-59* and *his-55*, using CRISPR/Cas9-mediated genome editing. Interestingly, removal of *his-59* and *his-55* is already sufficient to result in detectable phenotypes, including reduced brood size and increased germline apoptosis. We observed a variable penetrance of shrunken and collapsed germlines that exhibit a reduction in germ cell number. To understand whether this germ cell loss is due to a reduction of meiotic or/and mitotic cells, we used a reporter that specifically labels the mitotic cells by GFP (*41*), while all germ cells are labeled by mCherry. Comparing wild-type and *his-59*; *his-55* gonads, the progenitor mitotic zones were comparable, whereas the meiotic regions were greatly reduced in the double *H3* mutant (Fig. 3A and B). Consistently, *his-59*; *his-55* double mutants displayed significantly reduced brood sizes (Fig. 3C). To assess whether the germ cell loss and brood size reduction were due to increased cell death, we used a CED-1::GFP reporter to visualize apoptotic germ cells in adults (*42*). Increased germ cell apoptosis was detected in the *his-59* single *H3* mutant, and such a phenotype is enhanced in the *his-55; his-59* double mutant (Fig. 3D and E). Taken together, these data indicate that compromising H3 levels in the germline by removing two Class I *H3* genes causes increased germ cell apoptosis and germline atrophy. In contrast, previous studies demonstrate that H3.3 is dispensable, as knockout of multiple *H3.3* genes did not lead to any detectable germline defects, using a strain that lacks all H3.3 homologues, including *his-69, his-70, his-71, his-72*, and *his-74* (*29*). Taken together, these results indicate that H3 has an essential role in establishing or maintaining a unique chromatin state in the differentiating germline and this role cannot be compensated by H3.3.

**Fig. 3.**
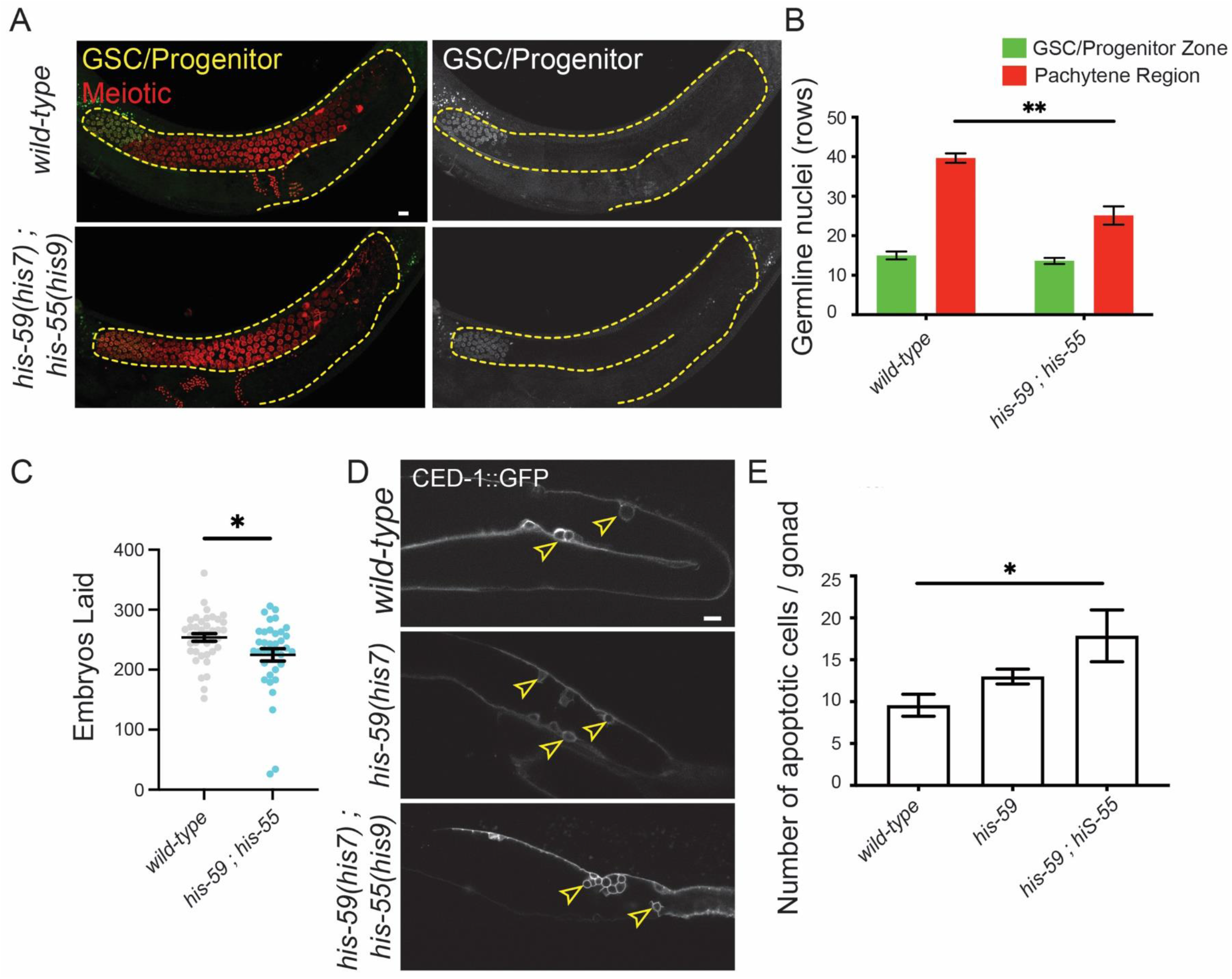
Knockouts of the germline-expressed histone *H3* genes lead to decreased fecundity and germ cell nuclei, as well as increased germline apoptosis. (**A**) Representative images of wild-type and *his-59(his7); his-55(his9*) double mutant of histone *H3* genes. The strain GC1413 *rrf-1*(*pk1417*; *naSi2* (*Pmex-5::H2B::mCherry::nos-2 3’UTR*); *teIs113* (*Ppie-1::GFP::H2B::zif-1 3’UTR*)) was used to label all germline nuclei with mCherry (red), while progenitor nuclei are doubly marked with GFP and mCherry (yellow). The dashed lines outline the gonads. (**B**) Quantification measured by rows of cells from the distal end. All quantifications = average ± SE. *P*-value: unpaired t test, showing no significant difference (*P* = 0.3192) in the Progenitor region (GFP- and mCherry-double positive germ cells) of *his-59(his7);his-55(his9*) double histone *H3* mutant (n=4) and wild-type (n=3), but a significant difference (\*\**P* = 0.0054) in the pachytene region (GFP-negative and mCherry-positive germ cells), *his-59(his7);his-55(his9*) double histone *H3* mutant (n=8) and wild-type (n=8). (**C**) Brood size of the *his-59(his7); his-55(his9*) double histone *H3* mutant (n=36) and wild-type (n=39) lines. Each data point represents the number of living larvae from individual worms with the corresponding genotype. (**D**) Representative images of nuclei undergoing programmed cell death marked by CED-1::GFP (yellow arrows). (**E**) Quantification of apoptotic cells per gonad of wild-type (n=7), *his-59(his7*) single mutant (n=5), and *his-59(his7); his-55(his9*) double mutant (n=7). All quantifications = average ± SE; *P*-value: unpaired t test, ** *P<0.01*, * P≤0.05. Scale bars:10 μm in (**A** and **D**).

Prior to gastrulation, embryonic blastomeres are characterized by decondensed chromatin and wide cellular differentiation capacity, termed as pluripotent (*43-45*). As embryos transit through gastrulation, they acquire distinct cell fates, accumulate heterochromatin, and restrict the developmental plasticity (*44, 46, 47*). To determine whether the increased incorporation of H3 during gastrulation is necessary for restricting embryonic plasticity, we generated a mutated form of H3 containing a single substitution of histidine (His, H) to aspartic acid (Asp, D) at the 113th position (H113D). The C-terminal H113 of H3 is a key residue that functions at the interface between the two H3-H4 dimers (H3 and H3’ in Fig. 4A). By changing the positively charged H to the negatively charged D, the H113D mutant is thought to drastically destabilize the H3:H3’ interface, preventing the complex formation between an H3-H4 dimer and the histone chaperone CAF-1. This mutant has been reported to have a dominant negative inhibition on CAF-1 mediated nucleosome deposition *in vitro* and *in vivo* (*48, 49*). In order to avoid detrimental developmental defects, we generated this mutation at the *his-6* locus, a somatic-specific Class II histone *H3* gene, which limits the expression of the H113D restrictively in the somatic lineage and specifically at the onset of gastrulation. The *his-6*(*H113D*) mutant worms were viable, with no apparent embryonic abnormalities or change in the timing of embryogenesis. We then examined whether H113D acts dominant negatively to reduce the deposition of wild-type H3. Indeed, when crossing the *his-6(H113D*) mutant into a reporter strain carrying both *his-55::GFP* (H3) and *his-72::mCherry* (H3.3), the *his-6*(*H113D*) resulted in a significantly decreased H3 incorporation during embryogenesis, when comparing to the wild-type strain at multiple timepoints in early embryonic development (Fig. 4B-C). By contrast, H3.3 incorporation showed either inconsistent (1^st^ and 2^nd^ histograms) or insignificant (3^rd^ and 4^th^ histograms) changes at the comparable embryonic developmental timepoints (Fig. 4D).

**Fig. 4.**
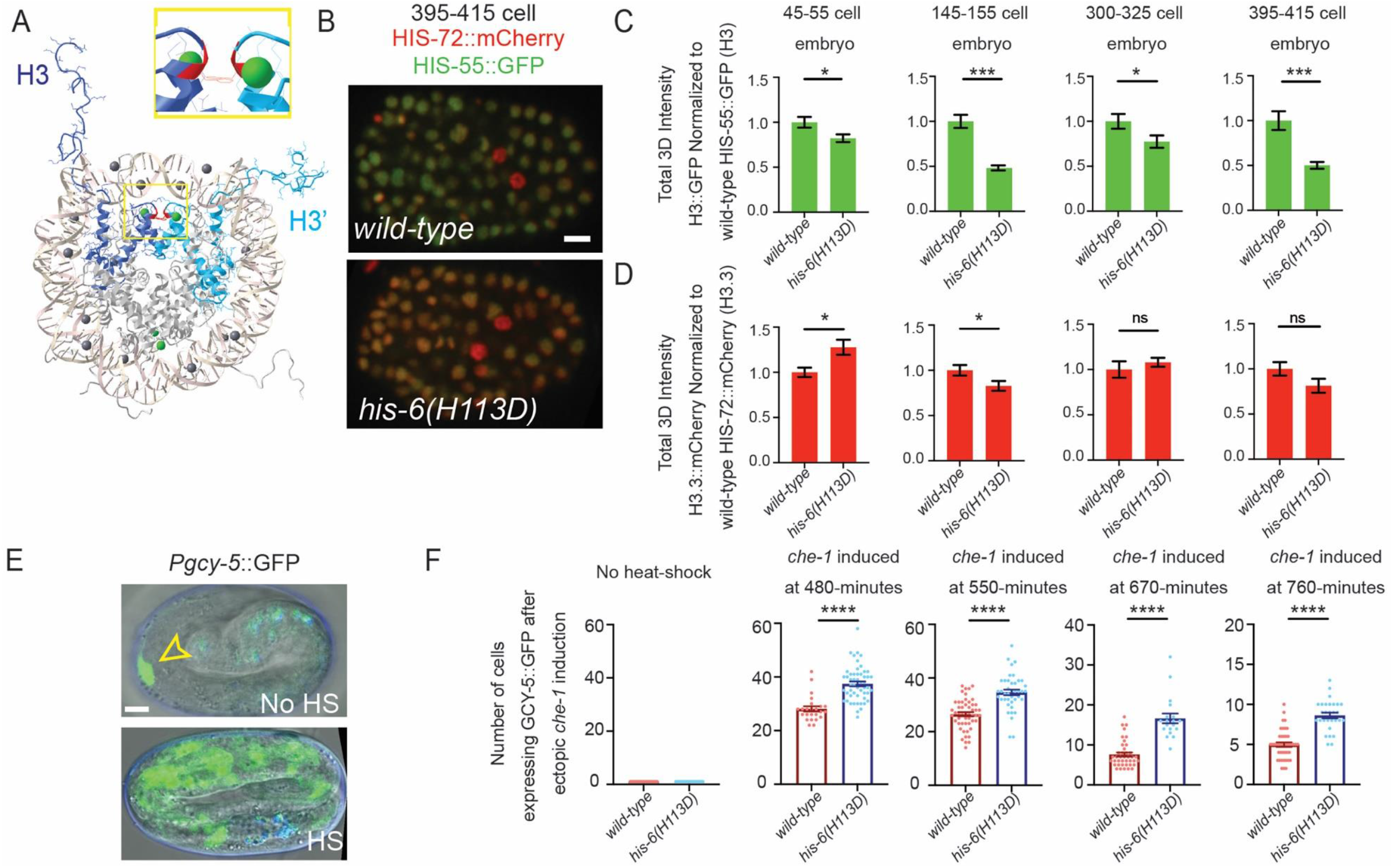
A dominant-negative mutation in the somatic-specific histone *H3* gene, *his-6(H113D*), destabilizes the histone tetramer (H3-H4)_2_, leading to reduced H3 incorporation and extended embryonic plasticity. (**A**) Structure of histone H3 in the histone octamer, drawn with Cn3D (https://www.ncbi.nlm.nih.gov/Structure/CN3D/cn3d.shtml). The two H3 in the octamer (H3 and H3’) are colored in dark blue (H3) and light blue (H3’). The interface of H3 and H3’ maintains the (H3:H4)_2_ tetramer, for which the H113 (Red) →D mutation in the somatic H3-encoding *his-6* gene has a dominant-negative effect by destabilizing the (H3:H4)_2_ tetramer specifically in somatic cells. A zoomed in view is displayed in the inset. (**B**) Representative images of embryos expressing H3.3::mCherry by *his-72::mCherry* and H3::GFP by *his-55::GFP* at the 395-415-cell stage to compare the H3::GFP levels in *wild-type* vs. *his-6(his10*)(*H113D*) mutant embryos. (**C-D**) Quantification of *his-72::mCherry (H3.3*) and *his-55::GFP (H3*) embryos at designated stages in *wild-type* vs. *his-6(his10*)(*H113D*) embryos. (**E**) Embryonic plasticity assay: The top panel displays an embryo without heat-shock induction of *che-1* expression, only the ASER neuron is labeled with bright *gcy-5::GFP* expression (yellow arrow) (signals that overlap with the blue channel are considered autofluorescence). The bottom panel is an example of heat-shock induced embryonic *che-1* expression at 550 minutes into embryogenesis, *gcy-*5 expression is broadly activated. (**F**) Quantification of the number of cells per embryo expressing *gcy-5::GFP* in response to *che-1* induction at different staged embryos. At each tested stage, embryos carrying the dominant negative mutation of *his-6(his10*)(*H113D*) display more *gcy-5::GFP*-expressing cells per embryo challenged during the same timepoints including 480-, 550-, 670-, and 760-minutes during embryogenesis, *his-6(H113D*) embryos resulted in a 38%, 33%, 118%, and 72% increase in cells that are converted, respectively. Scale bars: 5 μm. All quantifications = average ± SE; *P*-value: unpaired t test **** *P*≤*0.0001*, *** *P*≤*0.001*, ** *P*≤*0.01*, \**P*≤ *0.05*, ns: not significant.

The molecular processes underlying declined cellular plasticity during embryogenesis have been previously investigated (*44, 45, 50*). For example, all somatic lineages can be converted into other somatic cell types, depending on the ectopic expression of master cell fate defining transcription factors (*51-53*). However, this reprogramming flexibility is gradually lost in cells from embryos after initiating gastrulation (*44, 47, 51, 54-56*). It has been shown that ectopic expression of the transcription factor *che-1/*Zn-finger, which is normally expressed solely in the bilaterally symmetric ASE neuron pair (ASEL and ASER), is sufficient to induce the expression of a neuronal cell fate reporter (*gcy-5::GFP*) and aberrant adoption of neuronal cell fates in early embryonic blastomeres (*47, 57-60*). However, the ability of blastomeres to respond to ectopic expression of *che-1* is progressively lost during gastrulation (*47*). To determine whether the developmentally programmed increase of H3 deposition during gastrulation restricts the embryonic plasticity, we challenged embryos to adopt alternative cell fates at progressively later developmental timepoints. Without ectopic expression of *che-1*, the neuronal cell fate reporter, *gcy-5::GFP*, is expressed solely in the ASER neuron in late-stage embryos (Fig. 4E and F). In this assay, non-ASE cells that actively express *gcy-5::GFP* in response to *che-1* ectopic expression are considered plastic. When *che-1* is induced during early embryonic stages (480 and 550 min.) in a wild-type background, *gcy-5::GFP* is broadly activated (Fig. 4F). Consistent with the progressive loss of plasticity, *wild-type* embryos that are challenged later in embryogenesis (670 and 760 min.) demonstrate a reduced ability to respond to ectopically expressed *che-1* by turning on *gcy-5::GFP* (Fig. 4F). In contrast, in *his-6(H113D*) embryos, the number of cells that respond to *che-1* overexpression exhibited a significant increase in neuronal cell fate induction and *gcy-5::GFP* expression, when compared to *wild-type* controls throughout embryogenesis (Fig. 4F). Together, these results demonstrate that histone H3 incorporation during gastrulation progressively restricts embryonic plasticity and reduced H3 incorporation is sufficient to extend the window of embryonic plasticity.

Despite their widely divergent morphologies, invertebrates and vertebrates go through largely similar early embryonic stages. For example, it has been shown that the *de novo* establishment of heterochromatin domains during early embryogenesis is conserved from *Drosophila, C. elegans*, Zebrafish, to mammals (*46, 61-63*). Heterochromatin is known to be a potent epigenetic barrier against cellular reprogramming in *C. elegans*, as well as in mammals (*47, 64-66*). In *C. elegans*, the timing of heterochromatin formation during embryogenesis has been tracked using transmission electron microscopy and immunostaining with antibodies against heterochromatin-enriched PTMs (*46*). These studies have provided significant evidence that heterochromatin domains are established during gastrulation, coinciding with the drastic increase of *H3* gene expression and H3 incorporation identified in this work. Consistent with our results, extension of embryonic plasticity has also been identified in animals deficient in histone PTMs associated with heterochromatin, such as H3K9 and H3K27 methylation (*44, 47*). Furthermore, recent studies in *C. elegans* have identified that MET-2, an H3K9 methyltransferase, translocates from the cytosol to the nucleus during gastrulation. This changed localization and activity of the H3K9 methyltransferase may work together with the dynamic up-regulation of chromatin-associated H3 to co-regulate heterochromatin formation (*46*). Intriguingly, mouse early embryos transit from H3.3-enriched epigenetic landscape that is distributed evenly across the genome to a more canonical pattern (*67*). Therefore, our finding addresses the mechanism governing the developmental timing of *de novo* heterochromatin formation with its potential roles in antagonizing the cellular plasticity.

Histone PTMs have also been found to transit from both maternal and paternal chromatin to the zygote in *C. elegans*, as well as in mammals (*68-70*). For example, H3K27me3, which is generated by the Polycomb Repressive Complex 2, can be inherited both maternally and paternally (*69*). Therefore, histone PTMs established during oogenesis and spermatogenesis transmit an epigenetic memory from gametes to the zygote (*69*). Future investigations will be needed to determine whether unique histone PTMs are inherited preferentially on histone H3 or H3.3, as well as where H3 and H3.3 are distributed throughout the parental and early embryonic genomes in *C. elegans*.

In metazoans, histone genes are clustered together. For example, in the human genome, a large cluster on chromosome 6 and two small clusters on chromosome 1 contain all the RC histone genes. While in the mouse, a large cluster on chromosome 13 and two small clusters on chromosome 3 contain all RC histone genes (*11, 71*). In *C. elegans*, several clusters of histone genes are dispersed on four distinct chromosomes (Fig. S1). Here, our results demonstrate that the expression of each histone cluster is developmentally regulated for tightly coordinated expression, which is important for both germline fitness and embryonic development. Results from other metazoan embryos, such as sea urchin, suggest a potential conserved regulation of distinct histone gene clusters during embryogenesis (*72, 73*). Therefore, our results start to shed light on the functional roles of selective histone cluster expression during development. Given the conserved features of histone genes in different metazoan species, it will be interesting to investigate whether such a programmed histone gene expression is a common feature and what cue(s) may trigger such a co-expression.

## Acknowledgments

We thank Susan Strome, Geraldine Seydoux, John Kim, and members of the Chen Lab for helpful discussion of this work. We thank Johns Hopkins Integrated Imaging Center for confocal imaging; and Jane Hubbard for strains. Some strains were provided by the CGC, which is funded by NIH Office of Research Infrastructure Programs (P40 OD010440). We acknowledge support from NIGMS/NIH F32GM119347 (R.J.G), NICHD/NIH K99HD096053 (R.J.G.), NIGMS/NIH R35GM127075, HHMI Faculty Scholarship and Investigator program from Howard Hughes Medical Institute (X.C.).

## Author contributions

Conceptualization: RJG, XC

Methodology: RJG. XC

Investigation: RJG, CSS, GL

Visualization: RJG

Funding acquisition: RJG, XC

Project administration: RJG, XC

Supervision: RJG, XC

Writing – original draft: RJG, XC

Writing – review & editing: RJG, CSS, GL, XC

