## Supplemental Information for "Developmentally programmed epigenome regulates cellular plasticity at the parental-to-zygote transition"

**Supplementary Materials:**

**Materials and Methods:**

*C. elegans* strains

C. elegans were maintained on NGM (Nematode Growth Medium) agar plates using *Escherichia coli* OP50 as a food source and cultured according to standard methods (*74*). For heat-shock experiments, worms were grown at $15^0$C, heat-shocked at $32^0$C for 30 minutes, and left at $20^0$C overnight and scored about 24 hours later. Strains used and generated for this study are listed in Table S2.

CRISPR/Cas9 generated alleles

CRISPR engineering was generated by microinjecting Cas9-guide RNA (gRNA) ribonucleoprotein complexes into hermaphrodite gonads as described previously (*75*). Unique gRNA sequences were selected using the off-target predictions CRISPR design tool at https://crispr.mit.edu. For large edits, such as fluorescent protein tag sequences, we generated double-stranded DNA repair templates by amplifying eGFP, mCherry, or Dendra2 by PCR using specific oligos containing homology arms of approximately 35 bp. For histone H3 and H3.3, the fluorescent protein sequence for Dendra2, GFP, or mCherry was inserted into the C-terminus of the protein just before the stop codon. Tagging at the C-terminus was based on published H3.3 and H3 protein fusion analysis (*76*, *77*). For single-nucleotide modifications or deletions, we used single stranded oligonucleotides containing homology arms of approximately 35 bp as repair templates ordered from IDT, as standard 4 nM ultramer oligos. Silent mutations were included where necessary in the repair templates to modify either the PAM sequence or the gRNA seed region to prevent Cas9 from cleaving the repair template. (See table S3 for molecular details of gene editing reagents).

Immunocytochemistry

Germlines were dissected from 24-hour post-L4 adult hermaphrodites in egg buffer (25 mM Hepes, pH 7.4, 118mM NaCl, 48 mM KCl, 2 mM EDTA, 5 mM EGTA, 0.1% Tween-20, and 15 mM $NaN_3$) and fixed in 1% formaldehyde. Following fixation, samples were covered with a coverslip to ensure attachment to the slide surface, and flash-frozen on aluminum blocks chilled on dry ice. The samples were then fixed for 1 minute in 100% methanol (-20$^0$C) and rehydrated with PBST. Samples were then blocked for at least 30 minutes in PBS containing 0.1% BSA and 0.1% Tween 20 (PBST and BSA). Primary antibodies were diluted in PBST and BSA at the following concentrations: mouse anti-H3K27me2/3 diluted 1:5000 (39535, Active Motif), chicken anti-GFP diluted 1:250 (ab13970, abcam), and rabbit anti-H3K36me2 diluted 1:50 (ab9049, abcam). All primary antibody incubations were overnight at 4$^0$C. Slides were washed with PBST three times for 10 minutes each and incubated with secondary antibodies for 2 hours. Secondary antibodies were the Alexa Fluor-conjugated series (1:1000; Molecular Probes). Confocal images for immunostained fixed samples were taken using a Zeiss LSM 780 confocal microscope with 63x oil immersion objectives and processed using Fiji and Adobe Illustrator software.

Microscopy and Image analysis

For imaging experiments, many strains were surveyed under the microscope through the eye piece and general trends were noted for each strain generated. Then, a random subset of selected animals was imaged and analyzed more deeply, with quantitation. Live worms and embryos were mounted on 2% (wt/vol) agarose pads. Live-images were collected from 24-hour post-L4 adult hermaphrodites in M9 buffer and tetramisole (100mM) using a Zeiss 780 laser scanning confocal microscope. To stage embryos, two-cell embryos were selected and imaged using a Zeiss 780 laser scanning confocal, or a Zeiss spinning disk CSU-X1M for time-lapse imaging. Quantification of images were performed using the open-source Fiji software (*78*). Within any set of comparable images, the image capture and scaling conditions are identical. For each reporter, three randomly selected nuclear regions per animal were analyzed. To quantify the total amount of tagged-histone protein per nuclei, we conducted a 3D quantification by measuring the fluorescence signal in each plane from the Z-stack. Specifically, raw images as 2D Z stacks were saved as 16-bit TIF images, and the sum of the gray values of pixels in the image ("RawIntDen") was determined using Fiji. A circle was drawn to include all fluorescence signal (marked by Dendra2, GFP, or mCherry), and an identical circle was drawn outside the sample area as the background. The gray values of the fluorescence signal pixels for each Z stack were calculated by subtracting the gray values of the background signal from the gray values of the raw signal pixels. The total amount of fluorescence signal in the nuclei was then calculated by adding the gray values from all Z stacks. All quantifications of tagged-histone *H3* and *H3.3* samples were done using this method.

We analyzed the following number of embryonic nuclei in Fig. 2B: *his-72::Dendra2*: 2-cell (n=9), 4- cell (n=6), 20-50-cell (n=12), 115-175-cell (n=10), 230-300-cell (n=22) and 350-400-cell (n=21). *his-45::Dendra2*: 2-cell (n=11), 4- cell (n=29), 20-50-cell (n=9), 115-175-cell (n=18), 230-300-cell (n=16) and 350-400-cell (n=18). *his-6::Dendra2*: 20-50-cell (n=6), 115-175 cell (n=14), 230-300 cell (n=17) and 350-400-cell (n=18).

In Fig. 4C and D the following number of embryonic nuclei were analyzed in both wild-type and *his-6(H113D)* embryos: *wild-type*: *his-55::*eGFP(H3), *his-72*(H3.3)::mCherry at the 45-55-cell stage (n= 9,9), 145-155-cell stage (n=9,9), 300-325-cell stage (n=9,9), 395-415-cell stage (n=9,9). *his-6(H113D)*: *his-55::*eGFP(H3), *his-72*(H3.3)::mCherry at the 45-55-cell stage (n= 9,9), 145-155-cell stage (n=9,9), 300-325-cell stage (n=9,9), 395-415-cell stage (n=9,9).

<u>Analysis software</u>

Excel (Microsoft) and GraphPad Prism v9 were used for all data analysis and graphing. Images were captured with Zen Black (Zeiss) and Zen Blue (Zeiss). Fiji v1.53f was used for image processing and analysis. Photoshop (adobe) and Illustrator (adobe) were used for video editing and figure editing.

<u>Analysis of germline progenitor zone and pachytene nuclei</u>

The strain GC1413 *rrf-1*(*pk1417*; *naSi2* (*Pmex-5::H2B::mCherry::nos-2 3'UTR*); *teIs113* (*Ppie-1::GFP::H2B::zif-1 3'UTR*) was used to label all germline nuclei with mCherry (red), while progenitor zone nuclei are doubly marked with GFP and mCherry (yellow) in both wild-type and mutant backgrounds. Quantification of each region was measured by counting the rows of cells from the distal end (Figure 3A). For experiments where using this reporter was not possible due to overlap in the fluorescent channels, the germline regions were distinguished by row numbers based on the previously published analysis of the mitotic and meiotic regions (*79*, *80*). Specifically, the progenitor zone nuclei were selected for quantification from rows 1-5, the early pachytene nuclei were selected from rows 25-40, and the late pachytene nuclei were selected from rows 60-70 (Figure 1D).

Cell death assays

CED-1::GFP expressed in gonadal sheath cells, was used to count engulfed germ line corpses as described previously (*42*). Strains containing CED-1::GFP in wild-type and mutant backgrounds were maintained at $20^0$C. L4s of each genotype were selected and 24 hours later scored for the number of engulfed apoptotic cells in the gonad.

Brood size assays

Manually selected L4 animals were grown individually on petri dishes seeded with OP50 *E. coli* food until adulthood. They were then transferred on a new plate every 24 hours for a total of 4 days. The brood size of each worm was scored by counting the total number of larvae laid on the 4 plates. For each brood size experiment at least 30 worms were scored for each strain.

Cell fate challenge assay

Two-cell embryos were collected from wild-type and mutant backgrounds that carried the integrated reporter *otIs587 [gcy-5(fosmid::SL2::NLS::GFP + ttx3p::mCherry]* and *otIs304 [hsp16-2p::che-1::3xHA::BLRP + rol-6(su10060].* Embryos were incubated for 480 min, 550 min, 670 min, and 760 min. Heat-shock was administered at $32^0$C for 30 minutes, and left at $20^0$C overnight and scored about 24 hours later by counting the number of GFP positive cells. In this assay, non-ASE cells that activate expression of *gcy-5* in response to *che-1* ectopic expression are considered plastic. Images were acquired with a Zeiss LSM 780 laser scanning confocal microscope.

We analyzed the following number of wild-type and *his-6(H113D)* embryos in Fig. 4F: *wild-type*: No heat-shock (n=22), *che-1* induced at 480-minutes (n=24), *che-1* induced at 550-minutes (n=51), *che-1* induced at 670-minutes (n=40), *che-1* induced at 760-minutes (n=43). *His-6(H113D)*: No heat-shock (n=34), *che-1* induced at 480-minutes (n=52), *che-1* induced at 550-minutes (n=41), *che-1* induced at 670-minutes (n=19), *che-1* induced at 760-minutes (n=28).

**Legend for Supplementary Movie:**

**Supplemental Movie 1. Time-lapse movie of an embryo expressing H3.3 (HIS-72::mCherry) and a Class I H3 (HIS-55::GFP).** Live cell imaging of H3.3 (HIS-72::mCherry) and Class I H3 (HIS-55::GFP) during early embryogenesis. The dashed circles outline the P-lineage through multiple cell divisions. The video begins one frame prior to the first embryonic cell division and captures early embryogenesis until the P-lineage divides to generate the Z2/Z3 primordial germ cells. The video was acquired at 5-minute intervals. Snapshots are shown in Figure 2C.

### Figures and figure legends for supplementary materials

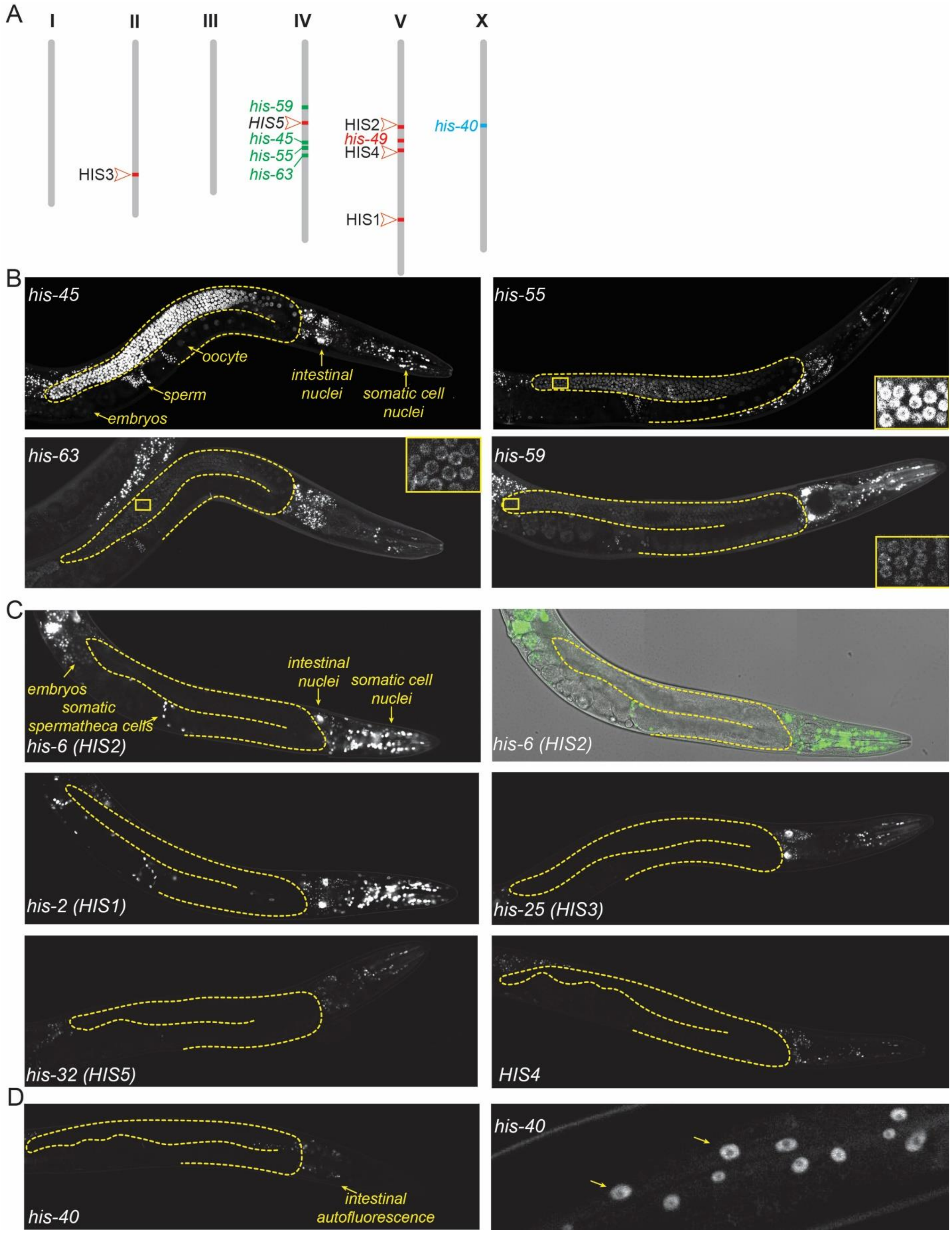

**Fig. S1. Expression patterns of all histone *H3* gene clusters in *C. elegans* adult hermaphrodites.** (**A**) Distribution of the different histone *H3* genes in the genome of the *C. elegans.* Bristol N2 strain is represented using a distinct color code for each class identified, including the ubiquitously expressed Class I histone *H3* genes (including both germline and somatic lineages) in green, somatic-specific Class II histone *H3* genes in red, and *his-40* in blue. (**B**) Representative fluorescence micrographs of ubiquitously expressed Class I histone *H3* isotypes including *his-45, his-55, his-63,* and *his-59*. The dashed lines outline the gonads, and distinct cell types are marked as an example. Insets demonstrate that *his-55, his-63,* and *his-59* are detectable in the germline by increasing the brightness. (**C**) Representative fluorescence micrographs for one member of each of the five histone gene clusters including HIS1 (*his-2*), HIS2 (*his-6*), HIS3 (*his-25*), HIS4 (*his-17*, *his-27*, and/or *his-49,* see methods for details), and HIS5 (*his-32*). Histone *H3* isotypes encoded in HIS1-5 are detectable in all somatic lineages, but undetectable in the germline. (**D**) *his-40* encodes a histone H3 that is detectable in epithelial nuclei including the hypodermis (epidermis) marked by yellow arrows.

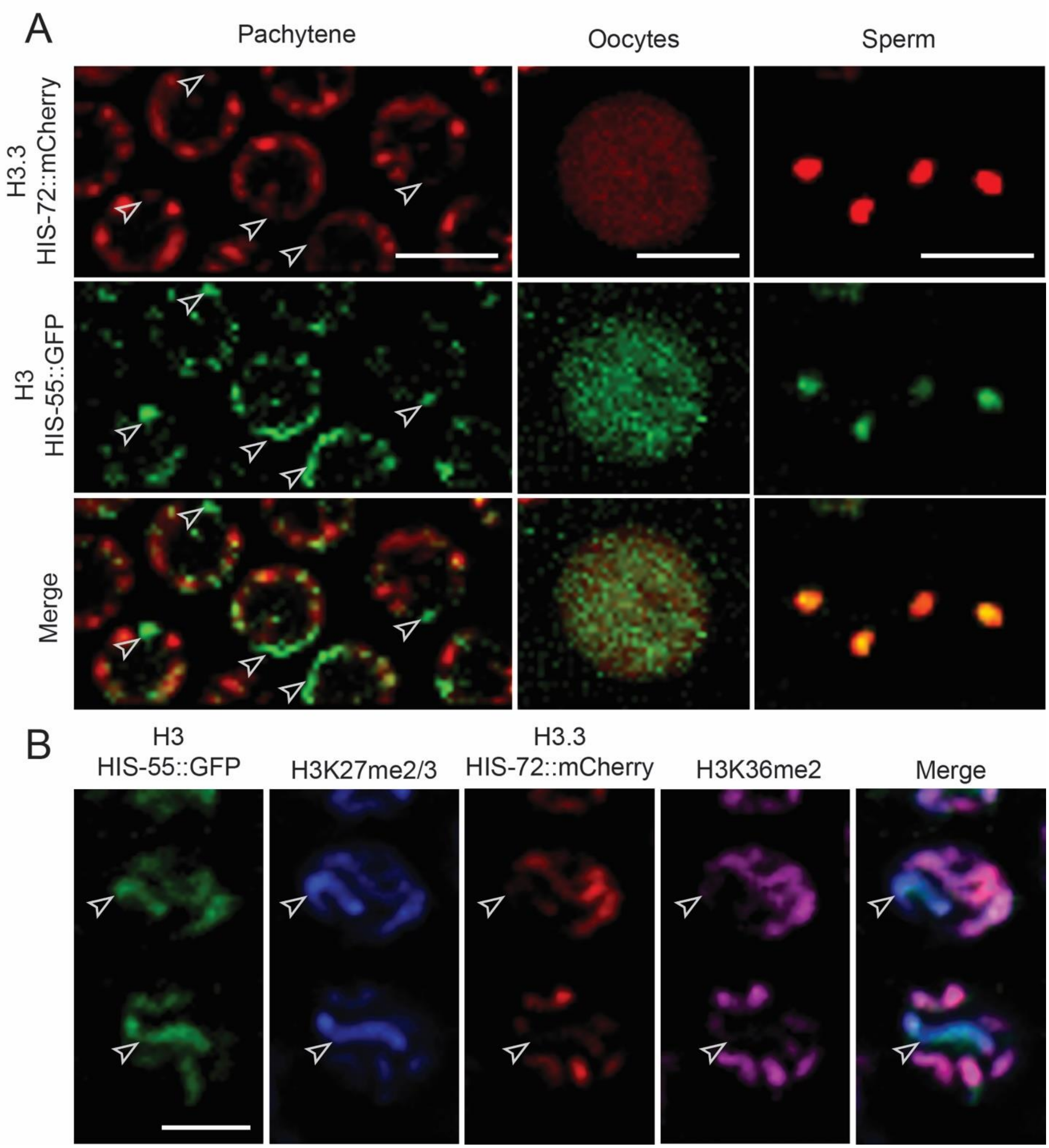

**Fig. S2. H3 and H3.3 occupy distinct chromatin domains in different staged germ cells.** (**A**) High magnification images of pachytene nuclei, oocyte, and sperm from a young adult hermaphrodite, demonstrating HIS-72::mCherry (H3.3) and HIS-55::GFP (H3). White arrows indicate chromatin regions showing presence of H3 but absence of H3.3. (**B**)

Immunofluorescence of HIS-72::mCherry (red), HIS-55::GFP (green), H3K27me2/3 (blue), and H3K36me2 (magenta) in pachytene nuclei. White arrows indicate Chromosome X identified by the presence of H3 and H3K27me2/3 but absence of H3.3 and H3K36me2. Scale bars: 5 μm.

**Supplemental Tables:**

**Table S1. Summary of histone H3-like, and histone H3.3-like expression in *C. elegans* hermaphrodites.**

| Gene (Chromosome) | Germ -line | Sperm | Oocyte | Pre-gastrulation | Gastrulation | Source |
|---|---|---|---|---|---|---|
| *his-45(H3) (IV)* | + | + | + | + | + | This study |
| *his-59(H3) (IV)* | + | + | + | + | + | This study |
| *his-63(H3) (IV)* | + | + | + | + | + | This study |
| *his-55(H3) (IV)* | + | + | + | + | + | This study |
| *his-2(H3) (V/HIS1)* | - | - | - | - | + | This study |
| *his-6(H3) (V/HIS2)* | - | - | - | - | + | This study |
| *his-9(H3) (II/HIS3)* | - | - | - | - | + | This study |
| *his-13(H3) (II/HIS3)* | - | - | - | - | + | This study |
| *his-17(H3) (V/HIS4)* | - | - | - | - | + | This study |
| *his-25(H3) (II/HIS3)* | - | - | - | - | + | This study |
| *his-27(H3) (V/HIS4)* | - | - | - | - | + | This study |
| *his-32(H3) (IV/HIS5)* | - | - | - | - | + | This study |
| *his-42(H3) (II/HIS3)* | - | - | - | - | + | This study |
| *his-49(H3) (V)* | - | - | - | - | + | This study |
| *his-40(H3) (X)* | - | - | - | - | + | This study |
| *his-72(H3.3) (III)* | + | + | + | + | + | Delaney et al., 2018 |
| *his-71(H3.3) (X)* | - | - | - | - | + | Delaney et al., 2018 |
| *his-69(H3.3-like) (III)* | - | - | - | - | - | Delaney et al., 2018 |
| *his-70(H3.3-like) (III)* | + | + | - | - | - | Delaney et al., 2018 |
| *his-74(H3.3-like) (V)* | + | + | + | + | PGC-restricted | Delaney et al., 2018 |

**Table S2.** ***C. elegans*** **strains used in this study.**

| Strain name | Genotype | Source | Comments |
|---|---|---|---|
| JHU42 | *his-45(his16[his-45::Dendra2]) IV* | This study | Dendra2 fusion |
| JHU5 | *his-6(his3[his-6::Dendra2]) V* | This study | Dendra2 fusion |
| JHU4 | *his-72(his2[his-72::Dendra2]) III* | This study | Dendra2 fusion |
| JHU20 | *his-72(his5[his-72::mCherry]) III* | This study | mCherry fusion |
| JHU19 | *his-55 (his11[his-55::TEV::eGFP::3xFlag]) IV* | This study | eGFP fusion |
| JHU14 | *his-59 (his7) IV* | This study | H3 homologue deletion |
| JHU16 | *his-59 (his7) IV; his-55 (his9) IV* | This study | H3 homologue deletion |
| JHU18 | *his-6 (his10) V* | This study | Histone H3 dominant negative *his-6(H113D)* |
| OH14454 | otIs587 [gcy-5(fosmid::SL2::NLS::GFP + ttx3p::mCherry]. otIs304 [hsp16-2p::che-1::3xHA::BLRP + rol-6(su10060] | CGC | GFP cell fate reporter |
| JHU45 | *his-6 (his10) V;* otIs587 [gcy-5(fosmid::SL2::NLS::GFP + ttx3p::mCherry]. otIs304 [hsp16-2p::che-1::3xHA::BLRP + rol-6(su10060] | This study | GFP cell fate reporter with histone H3 dominant negative allele |
| GC1413 | *Rrf-1(pk1417) I; naSi2(mex-5p::H2B::mCherry::nos-2 3'UTR) II; teIs113(pie-1p::GFP::H2B::zif-1 3'UTR) V* | Jane Hubbard lab (Roy D., et al., G3 2018) | Germline reporter |
| MD701 | *bcIs39 [lim-7p::ced-1::GFP + lin-15(+)]* | Zhou et al., 2001 | Apoptotic germ cell reporter |
| JHU57 | *his-55(his17[his-55::Dendra2]) IV* | This study | Dendra2 fusion |
| JHU59 | *his-63(his18[his-63::Dendra2]) IV* | This study | Dendra2 fusion |
| JHU53 | *his-59(his19[his-59::Dendra2]) IV* | This study | Dendra2 fusion |
| JHU39 | *his-2(his20[his-2::Dendra2]) V* | This study | Dendra2 fusion |
| JHU40 | *his-25(his21[his-25::Dendra2]) II* | This study | Dendra2 fusion |
| JHU37 | *his-13(his23[his-13::Dendra2]) II* | This study | Dendra2 fusion |
| JHU41 | *his-32(his22[his-32::Dendra2]) IV* | This study | Dendra2 fusion |
| JHU80 | *H3::Dendra2 in the HIS4 cluster (his-17, his-27, and/or his-49 ) V ** | This study | Dendra2 fusion |
| PHX2995 | *his-40(syb2995[his-40::Dendra2]) X* | This study | Dendra2 fusion |

*CRISPR/Cas9 Dendra2 knock-in reagents were designed to specifically edit a histone *H3* gene within the HIS4 histone cluster on chromosome V (see Figure S1A), which contains three histone *H3* genes *his-17*, *his-27*, and *his-49*. The genotyping results indicated that one or more of these three *H3* genes contains the *Dendra2* sequences, but their genomic DNA sequences are too similar to distinguish which one of the three contains the *Dendra2* sequence. Therefore, we have used this strain as an H3 reporter representing the entire HIS4 cluster activity.

### Table S3. Reagents used for generating CRISPR/Cas9 mediated fusion proteins, deletion strains, and point-mutations in *C. elegans*.

| Allele | sgRNA target sequences (PAM sites in bold) | Repair template homology arms** or repair template (deletion and point-mutations) |
|---|---|---|
| *his-45(his16[his-45::Dendra2]) IV* | GCGCGCTTAAATACCTTTT**TGG**<br>GCTTGCTCAACTACCAAAAA**AGG** | 5’gctaagcgagtcaccatcatgccaaaggata Tccaattggccagacgcatccgaggagagc gTgctcagcacgtgatgaacaccccgg gaattaacc<br>5’ ggccctaaagagggccgttgggttcggttaagtttt gagattaagcttActTaactaTcaaaaaaAgtatTtt accacacctggctgggcaggg |
| *his-6(his3[his-6::Dendra2]) V* | GGTGGGGGTTTGAATCGAAA**CGG**<br>ATCGAAACGGTCTCAAACTC**TGG** | 5’cgccaagcgagtcaccatcatgccaaaggacatcc aattggccagacgtatccgaggagaacgtgctcagca cgtgatgaacaccccgggaattaacc<br>5’ctaaagagggccgttgggttcggtgAgAgtttg aatTgaaacAgtTtcaaaTtctAgaaatcagaaa tttaccacacctggctgggcagg |
| *his-72(his2[his-72::Dendra2]) III* | AGTGCTTCGAGAATTCCTGA**TGG**<br>GAGCTTAAGCACGTTCTCCG**CGG** | 5’ccacgccaagcgcgtcaccatcatgccaaaggacat gcaactcgccagacgcatTcgTggagaGcgtgctca gcacgtgatgaacaccccgggaattaacc<br>5’ggaaaaatacgaggattatggtacaagttggattaaat gaatattaaaagtgcttTgagaattAgtAatgAagcttac cacacctggctgggcaggg |
| *his-72(his5[his-72::mCherry]) III* | AGTGCTTCGAGAATTCCTGA**TGG**<br>GAGCTTAAGCACGTTCTCCG**CGG** | 5’ccacgccaagcgcgtcaccatcatgccaaaggacatg caactcgccagacgcatTcgTggagaGcgtgctcag cacgtgatggtgagcaagggcgaggag<br>5’ggaaaaatacgaggattatggtacaagttggattaaatg aatattaaaagtgcttTgagaattAgtAatgAagcttactt gtacagctcgtccatg |
| *his-55(his11[his-55::TEV::eGFP::3xFlag]) IV* | CAATTGGCCAGACGCATCCG**AGG**<br>GCTTGCTCAACTACCAAAAA**AGG** | 5’ gcgagtcaccatcatgccaaaggatatccaattggccag GcgTatTcgGggagagcgcgctgagaacctctacttcca Aggag<br>5’gtggccctaaagagggccgttgggttcggttaagttttgag attaagcttgctTaactaTcaaGaaagAtatttacttgtcatc gtcatccttgtaatc |
| *his-55(his17[his-55::Dendra2]) IV* | CAATTGGCCAGACGCATCCG**AGG**<br>GCTTGCTCAACTACCAAAAA**AGG** | 5’gcgagtcaccatcatgccaaaggatatccaattggccagGc gTatTcgGggagagcgcgctcagcacgtgatgaacac cccgggaattaac<br>5’gtggccctaaagagggccgttgggttcggttaagttttgaga ttaagcttgctTaactaTcaaGaaagAtatttaccacacctgg ctgggcag |
| *his-63(his18[his-63::Dendra2]) IV* | GCGCGCTTAAATACCTTTTT**TGG**<br>GCTTGCTCAACTACCAAAAA**AGG** | 5’cgctaagcgagtcaccatcatgccaaaggatatccaattgg ccagacgtatccgaggagagcgtgctcagcacgtgatgaaca ccccgggaattaacc<br>5’ggccctaaagagggccgttgggttcggttaagttttgagatta agcttActTaactaTcaaaaaaAgtatttaccacacctggctg ggcaggg |

| | | |
|---|---|---|
| *his-59(his19[his-59::Dendra2]) IV* | GAGCGCGCTTAAATACCTTA**TGG**<br>AAGCTTACTTAACTACCATA**AGG** | 5'agtcaccattatgccaaaggatatccagctggccagacgtatccgaggagagcgcgctcagcacgtgatgaacacccc cgggaattaacctg<br>5'ggccctaaagagggccgttgggttcggtgagttttgagtt gaagcttacttaaTtaTTataagAtatttaccacacctggct gggcaggggg |
| *his-2(his20[his-2::Dendra2]) V* | CGGTGGGGTTTGAATTGAAA**CGG**<br>AAATTTAAGCACGTTCTCCT**CGG** | 5'gccaagcgagtcaccatcatgccaaaggacatccaattg gccagacgtatTcgCggagaGcgtgctcagcacgtgat gaacaccccgggaattaacc<br>5'ccctaaagagggccgttgggttcggtgAAgtttgaattA aaacgAtctcaaactttctgaaaatcagaaatttaccacacct ggctgggcagg |
| *his-25(his21[his-25::Dendra2]) II* | GCTGGCTCAGTACCATTGGA**AGG**<br>TCAAGCTGGCTCAGTACCAT**TGG** | 5'ctaagcgagttaccattatgccaaaggacatccaattggca Agacgtatccgaggagagcgtgctcagcacgtgatgaacac cccgggaattaacc<br>5'gtggccctaaagagggccgttgggttcggttagattttgag atcaagctgActTagtaTcattAgaagAcatTTAccaca cctggctgggcaggg |
| *his-32(his22[his-32::Dendra2]) IV* | AGCGTGCTTAAATGTCTTTG**TGG**<br>ACATTTAAGCACGCTCTCCT**CGG** | 5'cacgctaagcgagttaccatcatgccaaaggatatccagct ggccagacgcatTcgaggagaAcgtgctcagcacgtgatg aacaccccgggaattaacc<br>5'gtggccctaaagagggccgttgggttcggttatttgagatca agcttgtacaaaatatcTacaaagaTatttaccacacctggctg ggcagg |
| *his-13(his23[his-13::Dendra2]) II* | GCTGGCTCAGTACCATTGGA**AGG**<br>TCAAGCTGGCTCAGTACCAT**TGG** | 5'ctaagcgagttaccattatgccaaaggacatccaattggcaa Gacgtatccgaggagagcgtgctcagcacgtgatgaacaccc Cgggaattaacc<br>5'gtggccctaaagagggccgttgggttcggttagattttgaga tcaagctAgTtcagtaTcattAgaagAcatTTAccacac ctggctgggcaggg |
| *H3::Dendra2 in the HIS4 cluster (his-17, his-27, and/or his-49) V ** | ACTCTGAAAATCAGAAATTT**AGG**<br>AAATTTAGGCACGTTCTCCT**CGG** | 5'cacgccaagcgagtcaccatcatgccaaaggacatccaatt Ggccagacgtattcggggagagcgcgctcagcacgtgatgaa Caccccgggaattaac<br>5'gccctaaagagggccgttgggttcggtgggggtttgaatcga aacggtctcaaactctgaaaatTaAaGatttaccacacctggct gggcagg |
| *his-59 (his7) IV* | CCCACGGATTATCAACCTAA**AGG**<br>GAGCGCGCTTAAATACCTTA**TGG** | 5'ggcagccgttagtttcacttttctcacagtcccccaTA gattatcaaTctaaagAcaGCTACGataTcttatA gtagttaagtaagcttcaactcaaaactcaccgaacccaa cggccctc |
| *his-55 (his9) IV* | GATTATCAACCTAAACGCAA**TGG**<br>GCGCGCTTAAATACCTTTTT**TGG** | 5'ggcagccgttagtttcacttttctcacagtcccccaTA gattatcaaTctaaagAcaCCGGACTGTTAGT TGTTCAAAGGataTcttatAgtagttaagtaagct tcaactcaaaactcaccgaacccaacggccctc |
| *his-6 (his10) V* | GGTGGGGGTTTGAATCGAAA**CGG** | 5'gtcggactcttcgaggacaccaacttgtgcgcaatcGAC gccaagcgagtcaccatcatgccaaaggacatccaattgg ccagacgtatccgaggagaacgtgcttaaatttctgatttcT agaAtttGATATCgtttcAattcaaacTcTcaccgaa cccaacggccctc |

*CRISPR/Cas9 Dendra2 knock-in reagents were designed to specifically edit a histone *H3* gene within the HIS4 histone cluster on chromosome V (see Figure S1A), which contains three histone *H3* genes *his-17, his-27,* and *his-49*. The genotyping results indicated that one or more of these three *H3* genes contains the *Dendra2* sequences, but their genomic DNA sequences are too similar to distinguish which one of the three contains the *Dendra2* sequence. Therefore, we have used this strain as an H3 reporter representing the entire HIS4 cluster activity.

** for fluorescent protein tagging, Dendra2, mCherry, and eGFP were inserted into the endogenous locus to generate either H3 or H3.3 fusion proteins using a *dpy-10* co-CRISPR strategy (*75*). Custom crRNA sequences were designed to target the sgRNA target sequence listed. High-fidelity PCR was used to generate a linear repair template with 35 bp homology arms. Sanger sequencing was used to confirm the accuracy of the knock-in allele.
